# Computational assessment of transport distances in living skeletal muscle fibers studied in situ

**DOI:** 10.1101/2020.06.04.135566

**Authors:** Kenth-Arne Hansson, Andreas Våvang Solbrå, Kristian Gundersen, Jo Christiansen Bruusgaard

**Affiliations:** Department of Biosciences, University of Oslo, Oslo, Norway; Center for Integrative Neuroplasticity, Department of Biosciences, University of Oslo, Oslo Norway; Department of Health Sciences, Kristiania University College, Oslo, Norway; Department of Physics, University of Oslo, Oslo, Norway

## Abstract

Transport distances in skeletal muscle fibers are mitigated by these cells having multiple nuclei. We have studied mouse living slow (soleus) and fast (extensor digitorum longus) muscle fibers in situ and determined cellular dimensions and the positions of all the nuclei within fiber segments. We modelled the effect of placing nuclei optimally and randomly using the nuclei as the origin of a transportation network. It appeared that an equidistant positioning of nuclei minimizes transport distances along the surface for both muscles. In the soleus muscle however, which were richer in nuclei, positioning of nuclei to reduce transport distances to the cytoplasm were of less importance, and these fibers exhibit a pattern not statistically different from a random positioning of nuclei. Together, these results highlight the importance of spatially distribute nuclei to minimize transport distances to the surface when nuclear density is low, while it appears that the distribution are of less importance at higher nuclear densities.

## Introduction

The largest cells in the vertebrate body are the muscle fibers, and their vast size introduce logistical problems with respect to synthetic capacity and transport of macromolecules. For example, a human sartorius muscle has an average fiber length of 42 cm (Sandve *et al.*, 2011) and a fiber cross sectional area of about 2500 μm^2^ (Kann, 1957) which leads to a volume of 1050 nl. Most other mononucleated cells range 5-20 μm in diameter and, assuming a spherical shape, have volumes less than 0.004 nl. The cell body of an alpha motor neuron might have a diameter of 50 μm and assuming a spherical shape this gives a volume of about 0.07 nl. In addition, it might have an axon spanning 1 m, and with a diameter of 7 μm this yields a volume of 38 nl. The motor neuron might thus be the second largest cell type in a human body, albeit the sartorius cell still has a volume almost 30 times larger. When cells become larger, the average transport distances increase and thus influence the overall transport times. Hence, small and large cells operate under different time scales (Cadart *et al.*, 2019). In larger cells, molecules would acquire longer transport times to reach their target. Because diffusion times, *t*, scale to the distance travelled *L* as *t* ∝ *L*^2^, diffusion is more efficient at shorter distances. In contrast, active transport times scale linearly to the distance travelled, although the speed relies heavily upon the molecular kinetics of the motor proteins. Typically, a kinesin motor moves at a speed of 1μm/s along the microtubules (Pilling *et al.*, 2006), while the myosin V moves at 3μm/s on the actin filaments (Schott *et al.*, 2002).

Mammalian skeletal muscle cells are syncytia, and human muscle cells might have several thousand nuclei (Cristea *et al.*, 2010). The high number density of nuclei is believed to be required due to the large fiber volume and long transport distances, and both the positioning and the nuclear number seems to be regulated to overcome these challenges based on restricted synthetic capacity and the physical limitations to intracellular transport (Bruusgaard *et al.*, 2003; Bruusgaard *et al.*, 2006; Metzger *et al.*, 2012b; Gundersen, 2016). We have previously suggested that the myonuclei seems to be distributed as if to repel each other to minimize cytoplasmic transport distances in mammalian fibers (Bruusgaard *et al.*, 2003; Bruusgaard *et al.*, 2006). This optimization seems to be important in the fruit fly as wells (Manhart *et al.*, 2018), since perturbation of the nuclear positioning has been reported to impair muscle function (Metzger *et al.*, 2012a; Folker & Baylies, 2013).

The fact that these cells are multinucleated has led to the proposal of a so-called cytoplasmic-to-nucleus domain that signifies a theoretical cytoplasmic domain governed by a single nucleus (Strassburger, 1893). Each nucleus is surrounded by a synthetic machinery that seems to remain localized (Pavlath *et al.*, 1989; Mishra *et al.*, 2015), and it has been shown that some proteins are localized in proximity to site of expression both in vitro (Hall & Ralston, 1989; Pavlath *et al.*, 1989; Ralston & Hall, 1992a) and in vivo (Merlie & Sanes, 1985; Sanes *et al.*, 1991).

In hybrid myotubes in which one or a few nuclei were derived from myoblasts expressing non-muscle proteins it was found that mRNAs for nuclear, cytoplasmic and ER-targeted proteins had similar distributions, and were in most cases confined to distances 25-100 μm from the nuclei (Ralston & Hall, 1992b). Similar observations were made with virus infected isolated muscle fibers in culture, with the additional observation that virus coding mRNA did not venture into the fiber interior but remained around the nucleus at the fiber surface at least for the first 40 hours after infection (Nevalainen *et al.*, 2013). These and other observations support the notion that myonuclei support the expression of different genes within a compartmentalized region of a fiber (Cheek, 1985; Hall & Ralston, 1989; Pavlath *et al.*, 1989). The most prominent example is the acetylcholine receptor that is normally expressed only in the nuclei near the endplate, and expression outside this area (acetylcholine super-sensitivity) seems to require that the receptor is transcribed in nuclei along the entire length of the fiber (Merlie *et al.*, 1984; Gundersen & Merlie, 1993).

On the other hand, experiments where aggregated blastomeres from two different mouse strains expressing distinct forms of cytosolic metabolic enzymes display heterodimers of the enzyme in the syncytial muscle tissue (Mintz & Baker, 1967), and histological examination displayed no mosaicism in the cellular distribution of the enzyme variants in muscle fibers (Frair & Peterson, 1983). This means that some smaller proteins that are not trapped in structures such as synapses and sarcomeres can be distributed over longer distances, and that the nucleus of origin is of less importance.

To get a better understanding of how nuclear positioning may affect transport distances in muscle fibers we employed precise confocal imaging and 3D reconstruction of fibers labelled in vivo in the soleus and EDL muscle. These muscles differ in their myonuclear density and their nuclear distribution (Bruusgaard *et al.*, 2003), and thus potentially highlight variations in nuclear pattern due to differences in functional needs and composition of the interior. We therefore modelled each nucleus as an origin of transportation network, and derived the area and volume of responsibility by Voronoi segmentation (Du *et al.*, 2010) as well as transport distances to the surface and cytoplasm. Additionally, we used simulations of transport distances to analyze the energy trade-off by letting a protein either diffuse locally from a nucleus or being actively transported.

## Results

We have analyzed fiber segments of live surface fibers of EDL and soleus muscle in situ. Nuclei were generally confined to the fiber surface, hence we analyzed nuclear domains both in 2D (as surface domains along the sarcolemma) and in 3D (as volume domains of the fiber cytoplasm).

### Cross sectional area and nuclear number in individual muscle fibers from the EDL and soleus muscle

Quantification of nuclear number after injecting fluorescently labelled oligonucleotides, and fluorescent dextran, revealed that the EDL (Fig. 1A) fibers had approximately half as many nuclei (47±10 nuclei /mm) compared to the soleus fibers (88±29 nuclei/mm) (Fig. 1B). Reconstructions of fiber segments from confocal stacks (Fig. 1C and D) showed that the EDL fibers were about 31% larger than those in the soleus (Fig. 1F). On average, the EDL muscle had a CSA of 1091± 278.4 μm and the soleus 831.8± 241.3 μm^2^. These differences in nuclear number and size resulted in EDL fibers having volume domains 130% larger than soleus fibers (Fig. 1G), and 93% larger surface domains than in the soleus (Fig. 1H). The domain volumes in the EDL were on average 23468 ± 5930 μm^3^ and 10187 ± 3432 μm^3^ in the EDL and soleus muscle, respectively. In the EDL, surface domains were on average 2907 ± 582 μm^2^, while in the soleus they were on average 1508 ± 582 μm^2^.

**Figure. 1.**
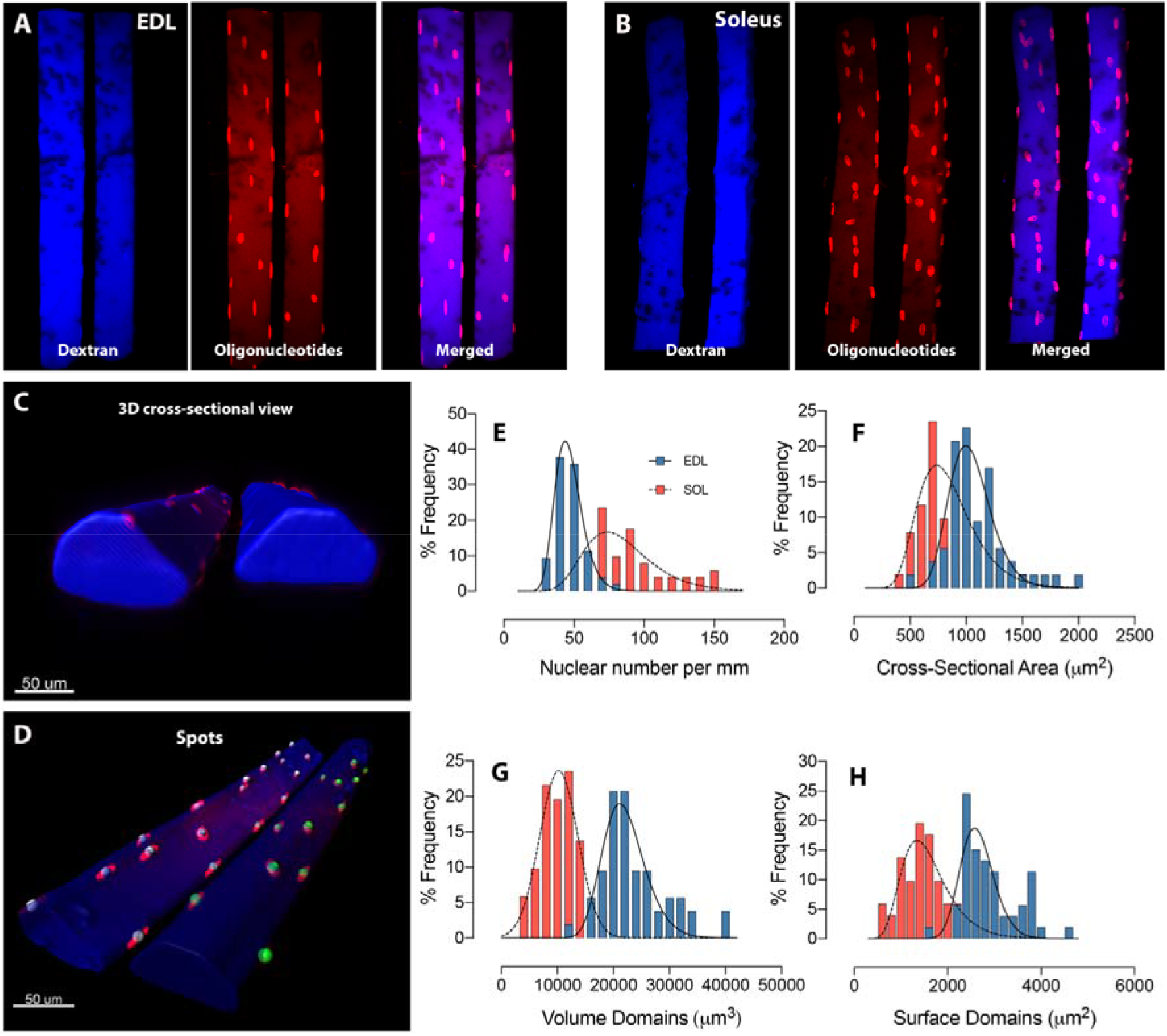
Three-dimensional modeling of muscle fibers after in vivo injections and confocal imaging. (A) Representative collapsed z-stacks of fibers from the EDL and (B) soleus muscles, injected with dextran (blue) and labelled oligonucleotides (red). (C) 3D modelled fibers based on the fluorescence from the cytosolic dextran. (D) Example of automatic assignment of spots to each myonucleus in order to define their 3D coordinates for subsequent analysis. (E) Frequency distribution of nuclear number and (F) cross sectional area, domain volume (G) and surface domains (H) in the EDL (blue) and soleus muscle (red).

### EDL and soleus display pattern differences of their domains

While the average nuclear domains are a simple function of the density of nuclei, and independent of how the nuclei are distributed, our data allowed us to study the distribution of the individual myonuclear domains. In a nuclear distribution where the distance between the nuclei is optimized, all domains would in principle be of equal size. Any deviation from that would imply a non-optimized positioning of nuclei, which may indicate that some domains would be unnecessarily large and represent “lacuna” where transport distances might be rather long.

To analyze how individual surface domains is influenced by the distribution of nuclei, we performed a detailed quantification of individual nuclear domains for each nucleus. Voronoi segmentation is a common way of formally quantifying areas or volumes of “influence” for distributed entities of the same kind. We modelled individual Voronoi domains by partitioning the fiber into sub-compartmentalized surface areas and volumes based on the position of each individual nucleus. To compare differences between nuclear distributions in the two muscles we used the standard deviation of the domain sizes. In EDL the observed values for the standard deviation measured for the Voronoi areas were not too different from those obtained when the nuclei were placed optimally, but some fibers showed much larger variability (Fig. 2B), indicating a less optimal distribution with some large lacunar domains. In the soleus, however, the observed pattern resembled more the values obtained when nuclei were placed randomly on the fiber surface (Fig. 2D). There was no obvious difference in the variation of nuclear position with fiber size (Fig. 2C and E).

**Figure 2.**
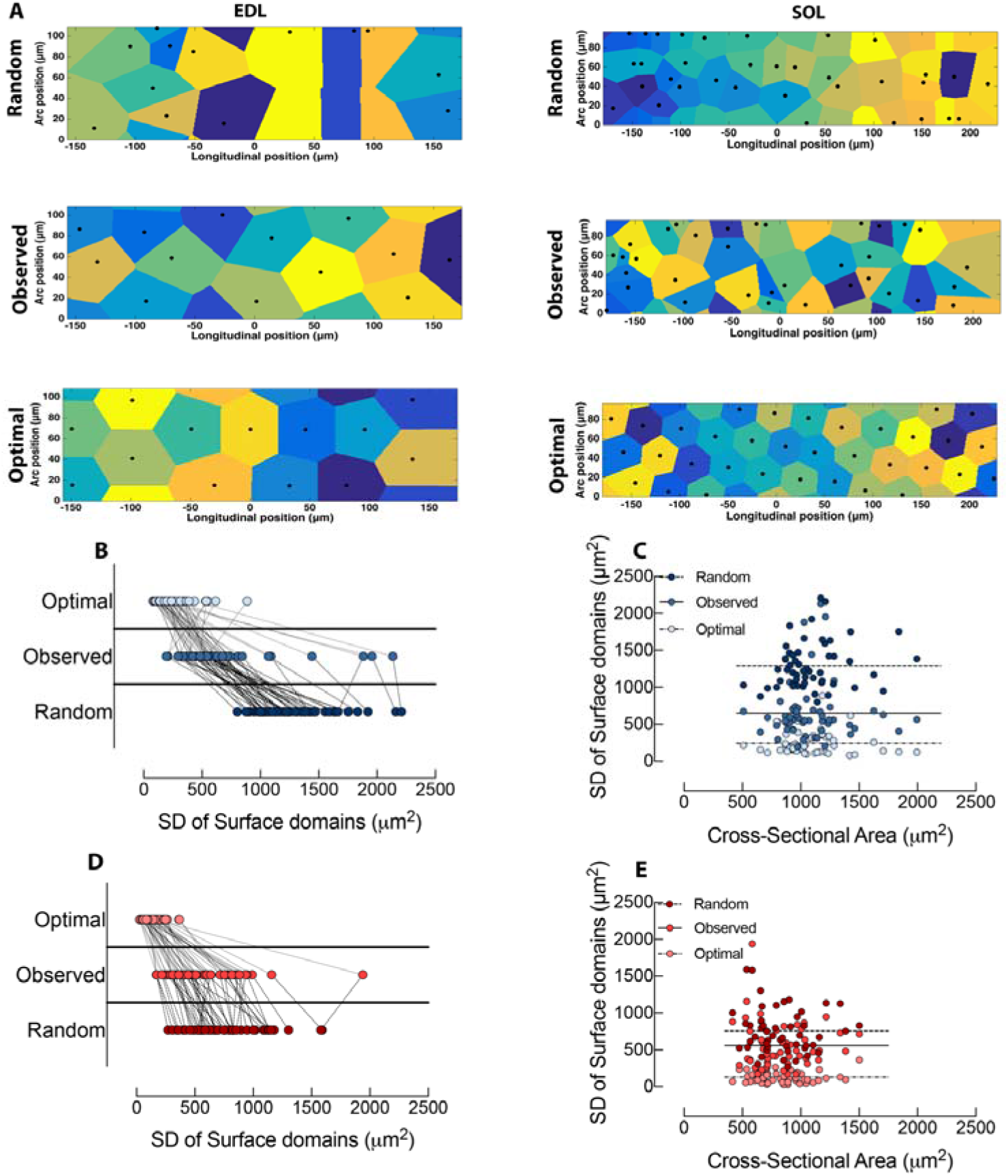
EDL and soleus display differences in patterning of Voronoi areas. (A) Representative Voronoi diagrams of individual domain areas in fibers from the EDL and soleus in the observed distribution and where nuclei were placed randomly or optimally on the fiber surface. (B) Comparison of observed and simulated nuclear patterns in the EDL and (D) soleus muscle plotted as a nested connection between the variation (standard deviation) of individual domains within a single fiber. Dashed and dotted lines signify the inter-specific connection between standardized variation of random-observed and observed-optimal. (C) Variation of surface domains plotted against the cross-sectional area in the EDL and (E) soleus. The relationship between variation and cross-sectional area was non-significant: Random (EDL, p=0.1625; SOL, p=0.5160), observed (EDL, p=0.8844; SOL, p=0.5858) and optimal (EDL, p=0.6892; SOL, p=0.9167).

Statistically, domain areas of the observed distribution were different from a random positioning of nuclei (mean difference in the SD of 641μm^2^; p<0.0001 in the EDL and 194μm^2^; p<0.0001 in the soleus) as well as optimal positioning (mean difference in the SD of, 405μm^2^; p<0.0001 for the EDL, and 432μm^2^; p<0.0001 for the soleus).

Next, we analyzed the Voronoi volumes (Fig. 3A), the difference between the two muscles resembled the values obtained when comparing Voronoi areas. Thus, in the EDL the variability resembled an optimal distribution (Fig. 3B), whereas in the soleus the variability was closer to placing the nuclei randomly (Fig. 3D). There was no clear effect of fiber size on the variation in Voronoi volumes (Fig. 3C and E).

**Figure 3.**
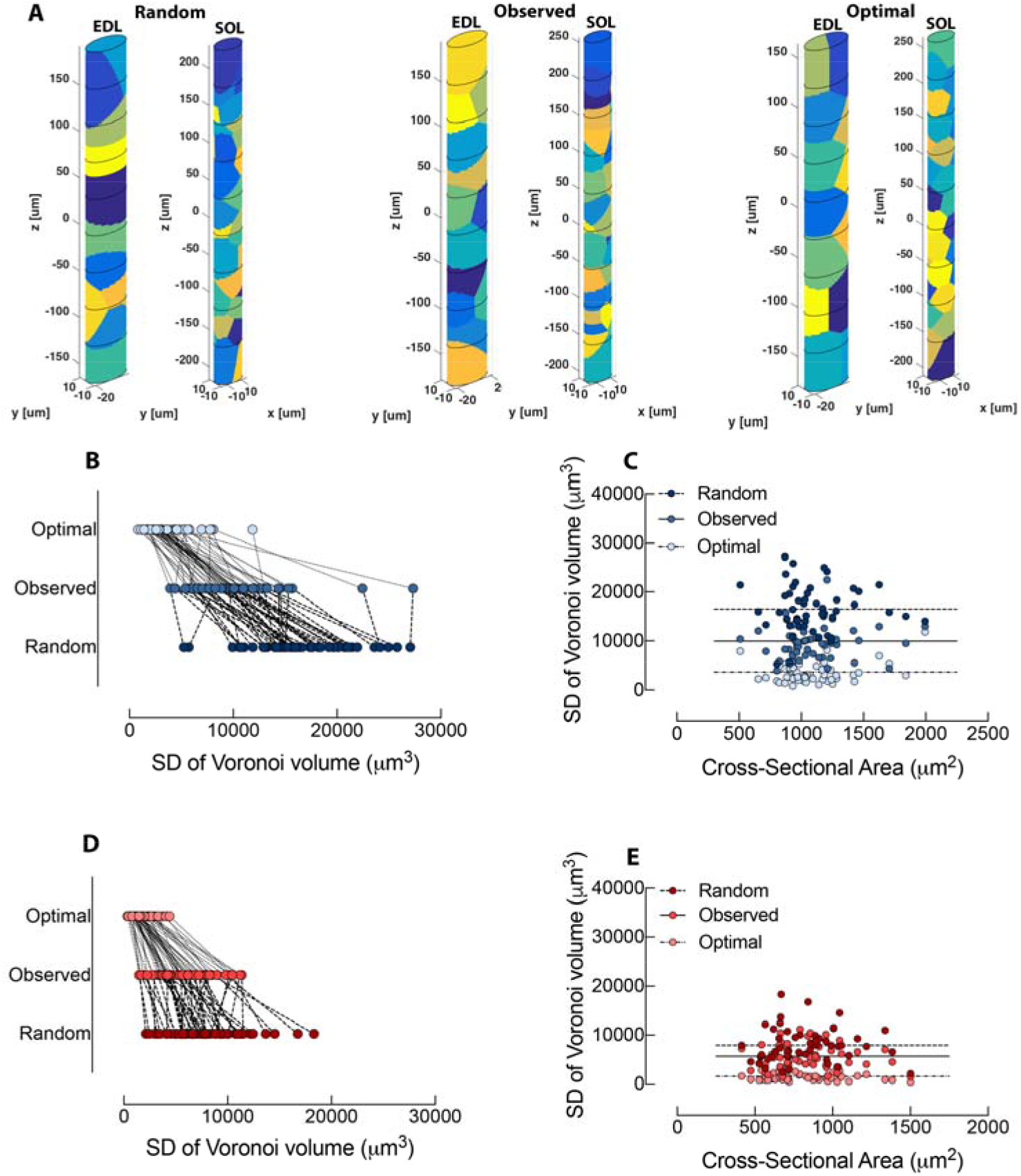
EDL and soleus display differences in patterning of Voronoi volumes. (A) Representative Voronoi diagrams of individual domain volumes in fibers from the EDL and soleus in the observed distribution and where nuclei were placed randomly or optimally on the fiber surface. (B) Comparison of observed and simulated nuclear patterns in the EDL muscle and (D) soleus plotted as a nested connection between the variation (standard deviation) of individual domains within a single fiber. Dashed and dotted lines signify the inter-specific connection between standardized variation of random-observed and observed-optimal. (C) Variation of surface domains plotted against the cross-sectional area in the EDL and (E) soleus. The relationship between variation and cross-sectional area was non-significant: Random (EDL, p=0.9651; SOL, p=0.9965), observed (EDL, p=0.9843; SOL, p=0.8789) and optimal (EDL, p=0.0627; SOL, p=0.4078).

Statistically, domain volumes of the observed distribution were different from a random positioning of nuclei (mean difference, 6521μm^3^; p<0.0001 in the EDL and mean difference, 224μm^3^; p<0.0001 in the soleus) as well as optimal positioning (mean difference, 6306μm^3^; p<0.0001 for the EDL, and 4084μm^3^; p<0.0001 for the soleus).

### Transport distances vary between the EDL and soleus muscle

The perhaps most relevant measure for judging the importance of nuclear density and positioning is the actual transport distances along the surface or through the cytoplasm. To model this, a grid of points corresponding to the pixel size was placed on the surface or in the volume of each fiber. The closest distance from these points to the nearest nuclei was calculated both on the surface (Fig. 4A) and in the fiber volume (Fig. 5A).

**Figure 4.**
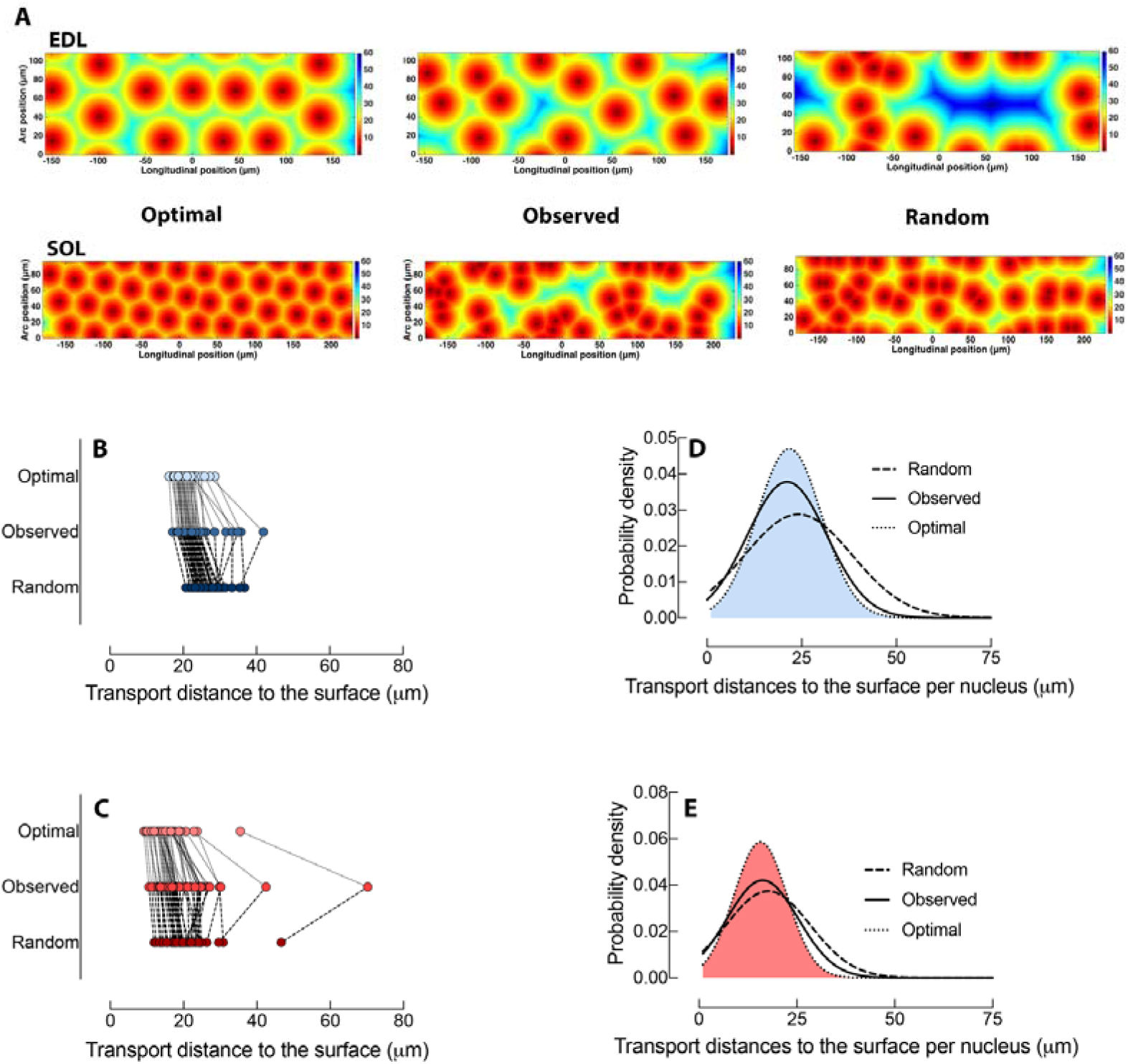
Quantitated measurements of transport distances along the fiber surface. (A) Fiber surface distance maps of fibers in the observed distribution and where nuclei were placed randomly or optimally on the fiber surface. (B-E) The average surface distance to the nearest nucleus were calculated and compared with optimal and random transport distances. (B) Transport distances in the EDL muscle (blue) compared by their mean value per fiber, and per nucleus after a Gaussian fit (D). (C) Transport distances in the soleus muscle (red) compared by their mean value per fiber and per nucleus after a Gaussian fit (E). Dashed and dotted lines in (B and C) signifies the inter-specific connection between random-observed and observed-optimal. The Red-Green-Blue (RGB) vertical scale bar/ color gradient in (A) signifies the distance at 0μm-30μm-60μm from its nuclear origin.

**Figure 5.**
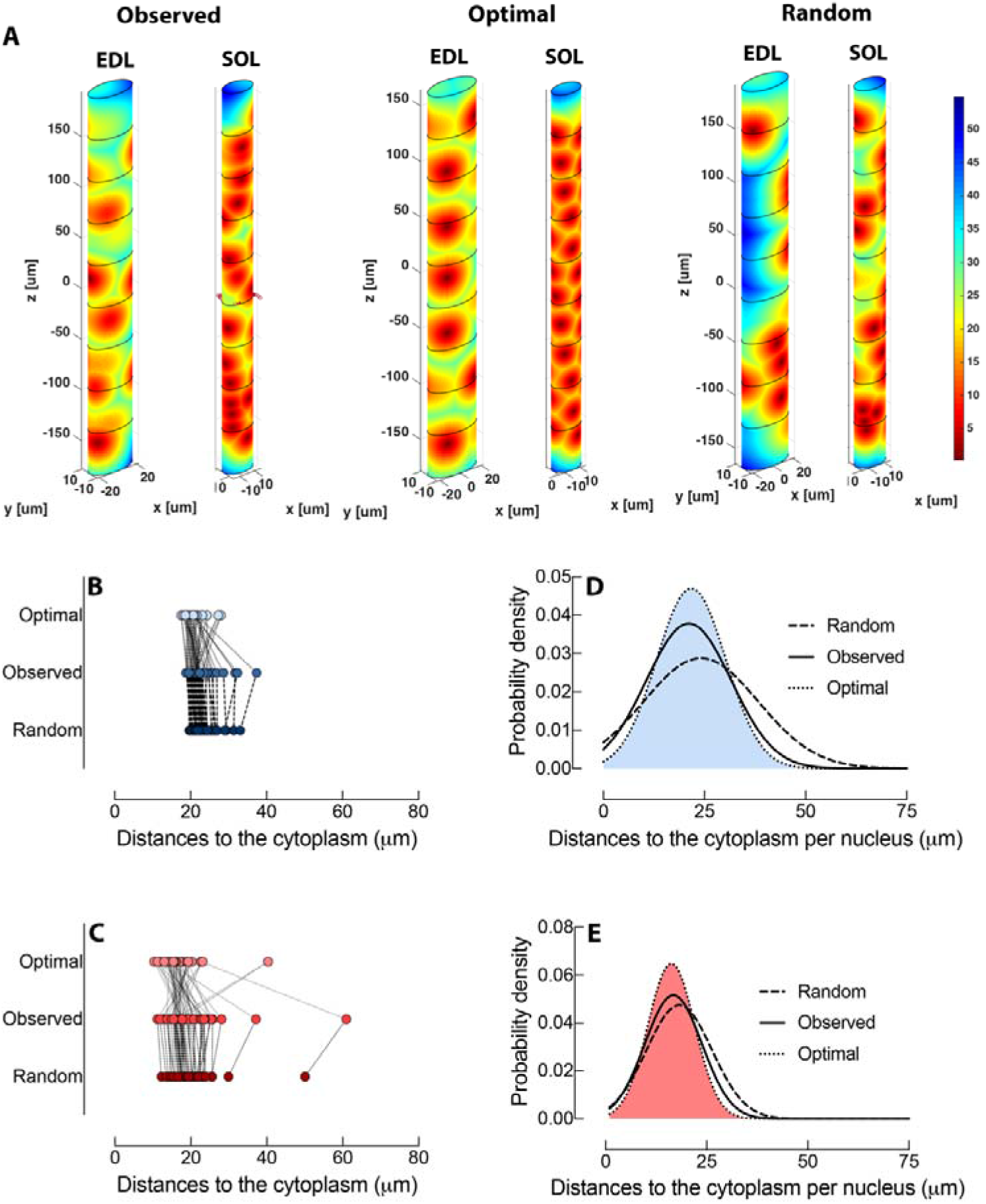
Measurements of transport distances to the cytoplasm. (A) Fiber volume distance maps of fibers where nuclei are placed optimally, randomly, and compared to the observed positioning. (B-E) The average surface distance to the nearest nucleus were calculated and compared with optimal and random transport distances. (B) Transport distances in the EDL muscle (blue) compared by their mean value per fiber and (D), per nucleus after a Gaussian fit. (C) Transport distances in the soleus muscle (red) compared by their mean value per fiber and (E) per nucleus after a Gaussian fit. Dashed and dotted lines in (B and C) signifies the inter-specific connection between random-observed, and observed-optimal. The Red-Green-Blue (RGB) color gradient in (A) signifies the distance at 0μm-30μm −60μm from its nuclear origin.

Comparing transport distances along the surface between different nuclear distributions showed that on average, EDL fibers had 10% longer transport distances (23± 4.7 μm) compared to the value obtained by placing nuclei optimally (21± 2.5 μm). When nuclei were placed randomly (27± 3.2 μm), the average distance were increased by 29% relative to optimal distances and the observed distances were closer to an optimal distribution (Figure 4B). In the soleus muscle, fibers had transport distances (20±9.2 μm) 25% larger compared to the average transport distance when nuclei were placed optimally (16±4.2 μm), while placing nuclei randomly (20±5.5 μm) was essentially identical to the observed distances (Fig. 4C).

For average transport distances in the cytoplasm (Fig. 5B), the observed distances in the EDL (22±3.6 μm) were closer to those seen when nuclei were placed randomly (23±2.7 μm) compared to nuclei placed optimally (21±2.6 μm). Interestingly, distances to the cytoplasm retrieved from placing nuclei randomly (20±5.5 μm) or optimally (17±5.7 μm) in the soleus (Fig. 5C) were not significantly different from observed transport distances (20±7.5μm).

In the EDL, there was a statistical improvement in placing nuclei optimally compared to the observed and random transport distances to the surface, and in the cytoplasm (Supp. Fig. 1A, B). However, there was no statistical difference between observed placement of nuclei and their transport distances to the cytoplasm when compared to optimally and randomly positioned nuclei (Supp. Fig.1B). In fibers of the soleus muscle, there was a statistical improvement when placing nuclei optimally for transport distances along the surface (Supp. Fig. 1C), but no improvement within the cytoplasm (Supp. Fig. 1D).

**Supplementary figure 1.**
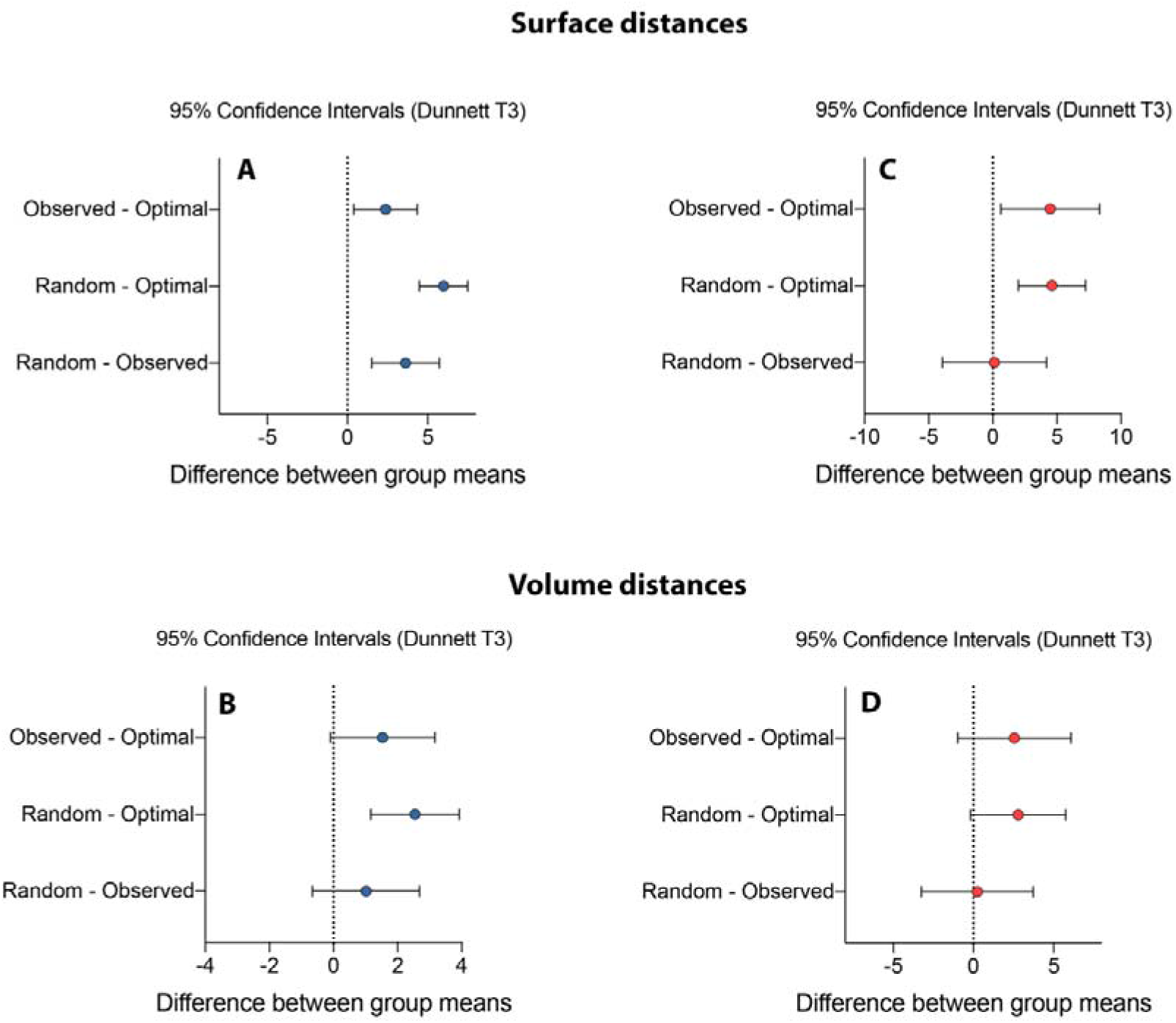
A group wise Brown-Forsythe analysis of differences in transport distances along the surface (EDL, A; SOL, C) and cytoplasm (EDL, B; SOL, D). For all graphs, 95% confidence intervals overlapping a zero difference between means is regarded as non-significant comparisons. A Dunnett post hoc test were performed to correct for multiple comparisons.

The average transport distance might not reflect whether there are some areas or volumes that is far away from any given nucleus, giving rise to lacunas that might have an insufficient supply of macromolecules. Therefore, we analyzed the distances on a per nucleus basis. Distribution of individual nuclear distances were fitted with a Gaussian function (Fig. 4D, E and Fig. 5D, E). It was evident from the Gaussian fit that the actual transport distances along the surface in the EDL where much improved compared to a random distribution and was approximating an optimal distribution (Fig. 4D). Similar conclusions were obtained analyzing optimal transport distances in the cytoplasm (Fig. 5D). Notably the longer transport distances were less prevalent by the observed placement of nuclei compared to a random distribution.

Soleus displayed similar traits when the transport distances along the surface (Fig. 4E) and cytoplasm (Fig. 5E) was analyzed, but the observed distribution was more similar to a random distribution.

In the EDL we found that 37% of the surface was longer than 25μm from the nearest nucleus, compared to 28% when nuclei were placed optimally and 47% randomly. In the soleus 27% of the fiber surface were longer than 25 μm from the nearest nucleus in the observed distribution, compared to 8% and 23% in the optimal and random distribution, respectively. Interestingly, and despite differences in fiber size and nuclear number, both muscles displayed about 97% of their surface to be within 50 μm of its nearest nucleus (Table I)

**TABLE I.**
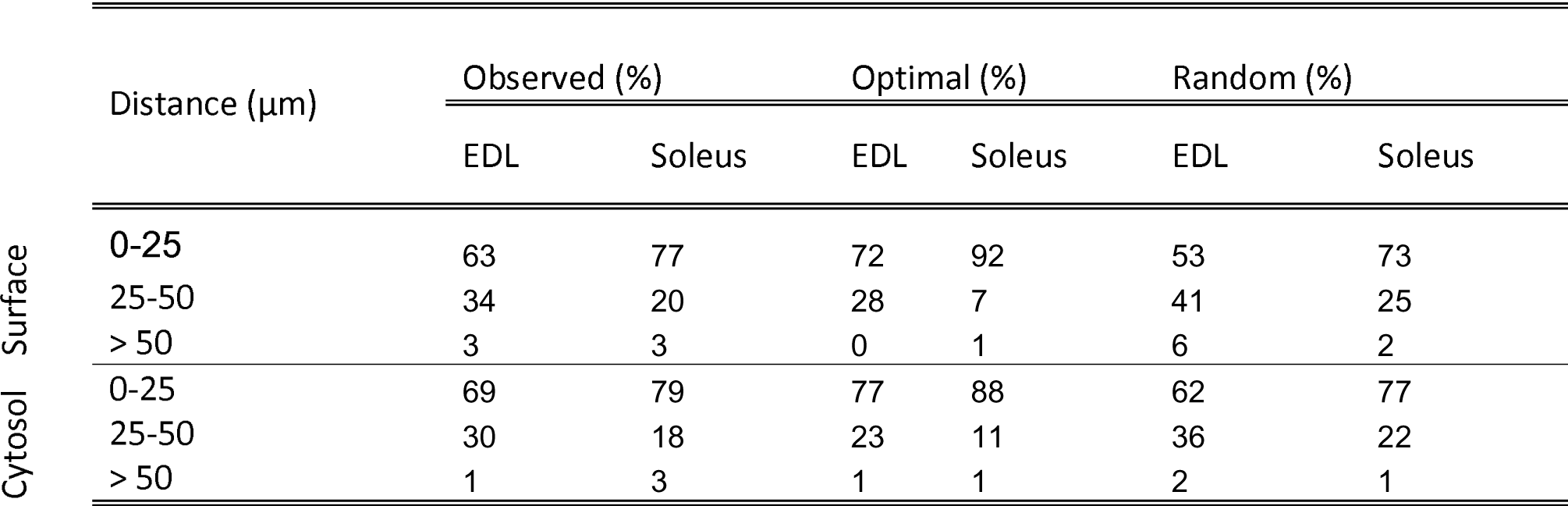
Percent of the cytosol or surface to its nearest nucleus at different distances.

In terms of volumes, 31% of the cytoplasm in the EDL were longer then 25 μm from its nearest nucleus during observed distributions, increasing to 38 % when nuclei were placed randomly, and 23% with optimal distribution of the nuclei. The soleus muscle displayed overall shorter transport distances as only 21% of the cytoplasm was more than 25 μm from its nearest nucleus, increasing only slightly to 23% when nuclei were randomly distributed, and 12% optimally. In spite of differences between the two muscles with respect to distances within 25 μm, 98% of the cytoplasm resides within a 50 μm proximity to the nearest nucleus.

### Consequences for diffusion times

We here describe the differences in dimensions, distribution and density of myonuclei in fibers from solus and EDL. It has however, previously been reported that also the effective diffusion coefficient differs in fibers from these two muscles (Papadopoulos *et al.*, 2000), and measured to be 12.5×10^−8^ cm^2^ s^−1^ and 18.7×10^−8^ cm^2^ s^−1^ in soleus and EDL respectively. When combined with our data, this information allows us to calculate diffusion times (*t*) from the nearest nuclei to any point in the cell according to *t* = *L^2^⁄2σD*, where *L* is the transport distance in *σ* dimensions, and *D* is the effective diffusion coefficient. The average diffusion times are given for each fiber in Fig. 6 and is compared to transport times that would be obtained by active transport (see Introduction).

**Figure 6.**
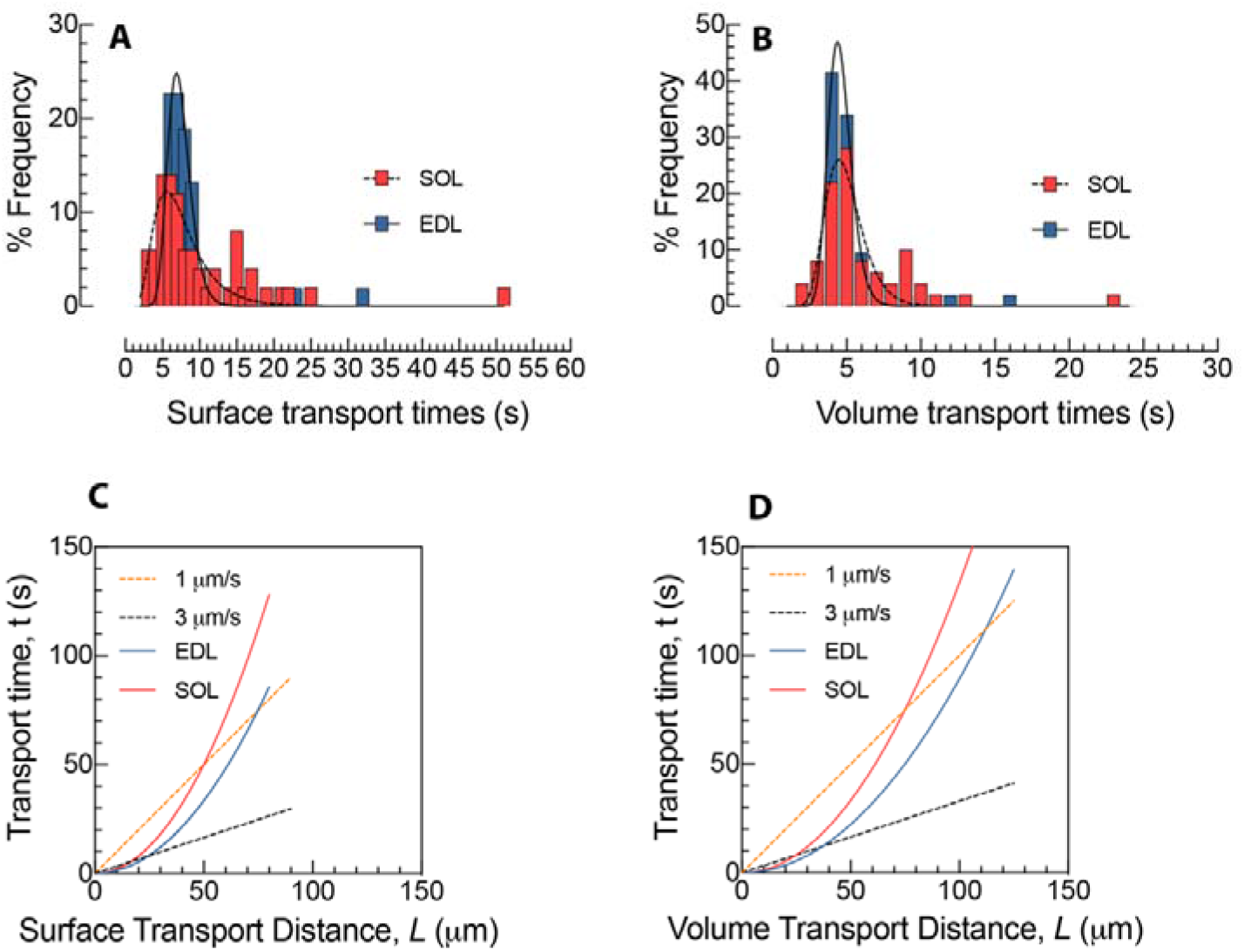
EDL and soleus muscle display roughly equal diffusion times. (A) Diffusion times plotted as the relative frequency distribution versus time along the surface and (B) within the fiber volume. (C) Transport times against distances along the surface and (D) within the volume Active transport at a speed of 3μm/s are highlighted by dashed lines. The intersection between dashed lines and the curvilinear data points highlight the interface where diffusion and active transport have equal transport times. At transport distances larger than the intersection, active transport is faster than diffusion.

We used myoglobin as an example and calculated the transportation times to each point on the surface. On average, we calculated transport times to be 8.2± 1.5s and 8.4± 1.8s for EDL and soleus respectively (Fig. 6A). For the diffusion times to each point in the entire cytoplasmic volume, the geometric mean was 5.0± 1.3s and 5.4± 1.6s for EDL and soleus respectively (Fig.6B). Thus, in spite of the large differences in dimensions, nuclear density and nuclear distribution, the diffusion time was remarkably similar for the two muscles.

It is likely that intracellular transport in muscle relies both on active and passive transport. Active transport along microtubules has been described in skeletal muscle (Pizon *et al.*, 2005; Wang *et al.*, 2013). While we are not aware of data on myosin mediated transport along actin in skeletal muscle, myosin IV has been observed localized to the fiber periphery, nuclei, sarcoplasmic reticulum and at the neuromuscular junction (Karolczak *et al.*, 2013), and the thick-filament-associated myosin molecule seems to be replaced in cultured myotubes in the absence of the microtubule system (Ojima *et al.*, 2015), while the actin network per se seems to be imperative for transport of membrane located molecules like e.g. the insulin sensitive glucose translocator GLUT4 (Brozinick *et al.*, 2004).

For the transport along the surface, diffusion would be faster than actin mediated transport (3μm/s) for distances of less than 15 μm in the soleus and 25 μm in the EDL. When compared to transport along microtubule, diffusion would be faster for distances up to 50 μm in the soleus and 75 μm in the EDL. (Fig. 6C).

When allowed to diffuse in three dimensions (Fig. 6D), diffusion would be faster than actin mediated transport for distances of less than 24 μm in the soleus and 36 μm in the EDL. When compared to transport along microtubule, diffusion would be faster for distances up to 75 μm in the soleus and 112 μm in the EDL.

## Discussion

We here present precise data for cell dimensions, and myonuclear density and positioning in fiber segments of living muscle fibers from EDL and soleus in situ. Fibers from these muscles varied with respect to all these variables, thus the radial size of the EDL fibers was 31% larger than those in the soleus, while the surface or volume per nuclei was 93% and 130% larger than in the soleus, respectively. In spite of these differences, the transport times to points on the surface or in the cytosol are remarkably similar in soleus and EDL, and we hypothesize that fibers are adapting the number and positioning of nuclei in order to achieve certain transport times for relevant macromolecules to all parts of the cell.

Given the differences in the variables related to cell dimensions and myonuclei it might seem like a paradox that the diffusion times are so similar, but is explained firstly by the diffusion coefficient being 50% higher in the EDL than in the soleus, secondly, we show that the nuclei are placed more optimally in the EDL. If the nuclei in the EDL were placed randomly, the average surface transport distance (∼27μm) versus the distance observed (∼23μm) would yield about 40% longer diffusion times, this would translate into a 30% lower concentration of a macromolecule at any given point, and the non-random positioning thus seems to be important in the EDL, and cytoplasmic dilution has been associated with impaired cell function in proliferative cells (Miettinen & Bjorklund, 2016; Neurohr *et al.*, 2019).

In contrast, in the soleus the placement of nuclei appeared close to random. It is possible that optimalisation of positioning is less critical since the density of nuclei is higher, another explanation is that other needs override those related to internal cell transport in oxidative muscles, thus in the rat soleus 81% of the nuclei appear next to blood vessels (Ralston *et al.*, 2006).

The interior of any cell is greatly crowded by macromolecules such as ribosomes, RNA and proteins (Zhou *et al.*, 2008; Smith *et al.*, 2017), which effectively reduce the motion of molecules and thereby impede diffusion. Additionally, muscle fibers have a dense mesh of myofibrils that occupy 80% of their total volume (van Ekeren *et al.*, 1992), which seems to act as exclusion zones for larger macromolecules (Papadopoulos *et al.*, 2000). Taking into account the effects of macromolecular crowding and scaling of transport times, it has been argued that mononuclear cells have small sizes in order to optimize diffusion and signaling (Soh *et al.*, 2013). They found the optimal size of a eukaryotic cell to have a radius of about 10 μm, about half of the transport distance found for muscle fibers in our calculations. This problem might be aggravated, since all nuclei do not express all genes at a given time-point (Fontaine & Changeux, 1989; Newlands *et al.*, 1998). Thus, the transport distances reported here might represent a bottleneck limiting cell function and size of muscle fibers.

Disturbance of myonuclear placement impairs muscle function e.g. after mutations in the Drosophila larvae (Metzger *et al.*, 2012b; Folker & Baylies, 2013). In mammals denervation, which leads to grave atrophy and is accompanied by myonuclear clustering eventually leaving lacunas with long transport distances (see introduction) (Viguie *et al.*, 1997; Ralston *et al.*, 1999). Similarly, nuclear distribution is disturbed in several myopathies and might play a causative role in impairing function (Romero, 2010).

## Material and Methods

### Animals

A total of 23 female NMRI mice of 20-30 grams of weight were used. The animal experiments were approved by the Norwegian Animal Research Authority and were conducted in accordance with the Norwegian Animal Welfare Act of 20th December 1974. The Norwegian Animal Research Authority provided governance to ensure that facilities and experiments were in accordance with the Act, National Regulations of January 15th 1996, and the European Convention for the Protection of Vertebrate Animals Used for Experimental and Other Scientific Purposes of March 18th, 1986.

### Preparing the mice for imaging

Before surgery, animals were anesthetized by a single intraperitoneal injection of a zrf cocktail (18.7 mg zolazepam, 18.7 mg tiletamine, 0.45 mg xylazine and 2.6 mg fentanyl per ml) that were administered at a dose of 0.08 ml per 20 g body weight.

The skin over the tibialis anterior and gastrocnemius was shaved and a small incision was made to expose the overlaying muscles that were subsequently retracted laterally to expose the EDL and soleus muscle, respectively. The epimysium was gently removed, and we took care not to damage the muscle. The exposed muscle was covered with a mouse Ringer’s solution; NaCl 154mM, KCl 5.6 mM, MgCl_2_ 2.2 mM and NaHCO_3_ 2.4 mM, and held in place with a coverslip mounted approximately 2 mm above the muscle. In some muscles, the neuromuscular end plate was visualized by applying Alexa 488-conjugated α-bungarotoxin (Molecular probes) to the surface of the muscle for 2–3 min.

In vivo intracellular injections of intravital dyes were essentially done as described previously (Utvik *et al.*, 1999). Animals were placed under a fixed-stage fluorescence microscope (Olympus BX50WI, Olympus, Japan) with a 20X, NA 0.3, long working distance water immersion objective. For *in vivo* labelling of nuclei and cytoplasm, single fibers in the EDL and soleus were injected with a solution containing 5 - TRITC or FITC-labelled random 17-mer oligonucleotide with a phosphorothioated backbone (Yorkshire Biosciences Ltd, Heslington, United Kingdom) and 2mg/ml Cascade blue dextran (10kDa, Molecular probes) dissolved in an injection buffer (10 mm NaCl, 10 mm Tris, pH 7.5, 0.1 mM EDTA and 100 mM potassium gluconate) at a final concentration of 0.5mM and 1mM, respectively. The oligonucleotides are taken up into the nuclei inside the injected fibers probably by an active transport mechanism (Hartig *et al.*, 1998), while dextran remain in the cytoplasm.

### Confocal Imaging

Muscle fibers and nuclei in the EDL and soleus were imaged with a confocal microscope (Olympus FluoView 1000, BX61W1, Olympus, Japan) in optical sections, separated by z-axis steps of 1 μm to have the full three-dimensional data set of nuclei. Sequential scanned fields of 0,09 mm^2^ were captured, with a pixel dwell time of 2μs. Movements induced by the mouse heartbeat or ventilation sometimes caused displacements of individual nuclei within stacks. Therefore, mice were given an overdose of the anaesthetics and image acquisition continued for 20 minutes after euthanasia. No difference in myonuclear positioning and fiber morphology was observed before and after euthanasia.

### Image analyzis

In vivo images (320 × 640 pixels × 2 μm voxel depth) from optical sections of muscle fibers were imported and analyzed for myonuclear number, 3D myonuclear positioning, fiber volume and surface area using the Imaris bitplane 8.3.1 software (Bitplane). Using the spot function in Imaris, a spot was automatically assigned to each nucleus based on the fluorescence intensity from the injected oligonucleotides, and if misaligned, spots were manually re-positioned to the center of each myonucleus. Highlighted spots/nuclei within fibers were then given a 3D coordinate in a relative Euclidean space using the Imaris Vantage extension. Volume and surface rendering were performed using the fluorescence from Cascade blue conjugated dextran in the cytoplasm. Rendering of fiber volume was performed using the fluorescence perimeter border of the fiber in the cross-sectional direction as an outer limit, and thereby preserving fiber morphology during quantification.

### Computer modelling of nuclear distribution

Once the Euclidian coordinates of the myonuclei were obtained in Imaris, fiber surfaces were approximated by fitting the points to the surface of an idealized elliptical cylinder surface.

An elliptical cylinder oriented along the z-axis, with its major and minor axes oriented along the x- and y-axes, respectively, can be parameterized as

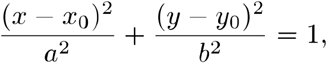

where *a* and *b* are the major and minor axes, respectively, and is the point where the center of the cylinder intercepts the *xy*-plane.

The cylinder was parameterized by the Euler angles pointing along the z-axis of the cylinder, the major and minor axes of the ellipse, the angle of the major axis relative to the x-axis, and the x- and y-coordinates of the center of the ellipse, for a total of 8 free variables. The optimization was carried out in two rounds, where the Euler angles were chosen first, after which the remaining ellipse fitting was completed for the given angles, using a direct least squares fitting. The angles were then updated using MATLABs interior point optimization solver, until convergence was achieved.

Subsequently, the 3-dimensional coordinates of the nuclei retrieved form Imaris were mapped to the 2-dimensional coordinates consisting of the position along the z-axis and distance along the ellipse circumference relative to the major axis. Next, we calculated area and volumes defined by Voronoi geometries of each nucleus and distance maps for all points on the cylinder surface to the nearest nuclei, based on the distance along the cylinder surface, noting that the surface is periodic along the circumference.

Additionally, we ran Monte Carlo simulations and nuclei were randomly placed on the parameterized surface in order to compare the distance map to that of the observed fibers. By a Lloyd’s Algorithm, we also placed the nuclei optimally on the surface, such that the Voronoi domains for all nuclei were equal and compared it to an observed distribution of nuclear patterns.

All codes used in the computational modelling can be found at https://github.com/CINPLA/cylinderTools.

### Statistics

Data were derived from 53 fibers injected on the lateral surface of EDL (N=9 animals) and 51 fibers injected at dorsal surface of the soleus muscle (N=12 animals). Descriptive statistics are shown as the pooled sum of fibers for a given muscle. If not stated otherwise, numbers are presented as mean± standard deviation (mean± SD). Frequently, point pattern analysis, including Voronoi segmentation, subdivide the area or volume of interest into a subset of polygons (2D) or polyhedrons (3D). In Voronoi segmentation, the number of Voronoi polygons is proportional to number of points in the plane or room. In our case, the number of points is equal to the nuclear number within each muscle fiber. Thus, if an area (or volume) is divided into the same number of domain objects, the average size of objects remains the same in spite of differences in the nuclear patterning (i.e. because the total area, volume and number of points are unchanged). Instead, we used the variation (standard deviation) as a measure of similarity between the observed mean size of polygons, to those when nuclei were placed randomly or optimally. We statistically compared standard deviations of domain areas and domain volumes by differences in nuclear patterning by a one-way ANOVA followed by a Tukey’s correction for multiple comparisons.

In comparison, we are not faced with the same problem as outlined for the Voronoi segmentation when analyzing transport distances between different nuclear distributions. They were therefore analyzed statistically by a Brown-Forsythe analysis followed by a Games-Howell’s multiple comparisons test at a 5 % significance level.

All graphs and statistical analyzis were plotted and performed in Prism 8 (Graphpad).

## Notes

### Competing Interest Statement

The authors have declared no competing interest.

